# Modulation of Electrotonic Coupling in the Inferior Olive by Inhibitory and Excitatory Inputs: Integration in the Glomerulus

**DOI:** 10.1101/072041

**Authors:** Jornt R. De Gruijl, Piotr A. Sokol, Mario Negrello, Cris I. De Zeeuw

## Abstract

Dendritic spines in glomeruli of the inferior olive are coupled by gap junctions and receive both inhibitory and excitatory inputs, yet the function of this configuration remains to be elucidated. In this issue of *Neuron*, Lefler et al. (2014) show that the GABAergic input from the cerebellar nuclei to neurons of the inferior olive dynamically affects both their coupling and subthreshold oscillations for short periods, while Mathy et al. (2014) and Turecek et al. (2014) show that the glutamatergic input to the olivary neurons can instruct plasticity of coupled dendrites in the long-term. The data sets from all three papers can be reconciled when put together in a model of the neuropil comprising the characteristic olivary glomeruli.

---

Whereas the intense debate between Santiago Ramón y Cajal and Camillo Golgi on the neuronal versus reticular doctrine of the central nervous system has been resolved more than a century ago, potential interactions between electrotonically coupled neuronal networks and afferent systems involving chemical neurotransmission have been hardly addressed in the mammalian brain (Spira et al., 1976; Landisman and Connors, 2005). The inferior olive probably forms one of the best nuclei to study these interactions, because it contains the highest density of neuronal gap junctions (De Zeeuw et al., 1998), the vast majority of which is situated between peripheral dendritic spines with extremely long necks in glomeruli that receive prominent afferent inputs (Sotelo et al., 1974; De Zeeuw et al., 1990). More specifically, most olivary spines that are coupled by gap junctions receive both an inhibitory synaptic input from a particular nucleus in the hindbrain and an excitatory input from either an ascending or descending projection from spinal cord or brainstem (Figure 1) (De Zeeuw et al., 1998). Interestingly, olivary neurons are also endowed with a set of conductances differentially distributed across the dendritic and somatic membranes that allow them to oscillate at subthreshold level, providing a means to generate ensembles of neurons oscillating and firing in synchrony (Llinás and Yarom, 1981). Thus, if one can decipher how chemical synapses in the inferior olive modulate the efficacy of electrical synapses, one may understand how the climbing fibers derived from the olive trigger spatiotemporal patterns of complex spike activity in Purkinje cells of the cerebellar cortex and thereby influence motor performance in the short-term and motor learning in the long-term (Van der Giessen et al., 2008).

In this issue of *Neuron*, Lefler et al. (2014) elegantly show with the use of optogenetic stimulation that the GABAergic input from the cerebellar nuclei to the inferior olive can cause a direct but transient decrement in electrical coupling between olivary cells (Figure 1). This effect can most likely be attributed to stimulating the input to the distal, dendritic and spiny synapses, because the authors measured, apart from a prominent impact on the coupling, a relatively consistent level of somatic postsynaptic responses to illuminations of different parts of the peripheral dendritic tree. The waveform of these post-synaptic potentials showed the characteristic frequency-dependent asynchronous release found following electrical stimulation of nucleo-olivary afferents (Best and Regehr, 2009). Based on this response the authors designed a stimulation pattern that elicited a prolonged, smooth shunt in the distal dendrites of the inferior olive compatible with the glomerular shunting hypothesis advocated by Llinás (1974). Interestingly, the authors also report that the strength of the coupling between two olivary neurons is asymmetric and that the level and even direction of this asymmetry can be regulated by stimulating their GABAergic input, highlighting the possibility that the spatial configuration of the patterns of complex spike activity can be created in a flexible fashion next to the temporal aspects. This finding is in line with the asymmetric distribution of cerebellar GABAergic terminals within the mammalian olivary glomerulus (De Zeeuw et al., 1990) as well as with the concept of asymmetric coupling mediated by network properties developed for Navanax (Spira et al., 1976). Thus, even though the spatial resolution of optogenetic stimulation and electrophysiological recordings has not yet reached the level of individual glomeruli and the concept of (asymmetrically) shunting coupled dendrites within glomeruli by chemical transmission remains to be demonstrated unequivocally, the data provided by Lefler and colleagues go a long way. The authors also show that activation of the GABAergic cerebellar fibers temporarily abolishes subthreshold oscillations, again without necessarily changing the somatic membrane potential. According to the authors, this result corroborates the idea that oscillations are in part an emergent network phenomenon, which depends on gap junctional current flows. This explanation follows the finding that the majority of GABAergic synapses are on the dendritic spines connected by gap junctions, with far fewer synapses located directly on the dendritic shaft or soma (De Zeeuw et al., 1998). Moreover, the network character of their findings on suppression of oscillations due to GABAergic input is also in line with the increased excitability of olivary cells when GABAergic modulation of coupling within olivary glomeruli is abolished. For instance, acute stimulation of excitatory afferents increases doublet firing in both connexin36-deficient animals, in which olivary coupling is absent, and mammals suffering from olivary hypertrophy, in which GABAergic input to the olive is affected (Ruigrok et al., 1990; Van der Giessen et al., 2008).

**Figure 1.**
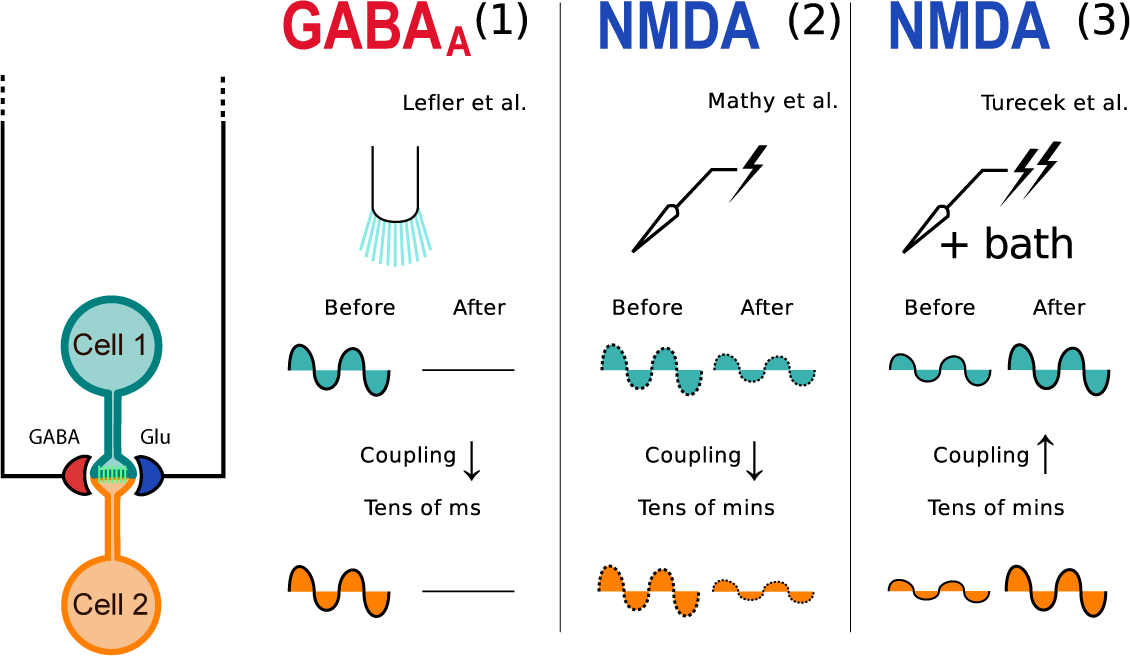
Experiments on gap junction modulation demonstrate multiple possible forms of plasticity. Lefler et al. (1) show that optogenetic induction of GABA release at the glomerulus creates a transient reduction of coupling coefficient; Mathy et al. (2) demonstrate that stimulation of pairs of inferior olivary cells at lower frequencies decreases the coupling coefficient; and Turecek et al. (3) reveal that stimulating olivary efferents to the glomerulus at higher frequencies increases the coupling coefficient. The waveforms in the diagrams represent the actual changes in subthreshold oscillation amplitudes as observed in (1) and (3), and the presumptive changes extrapolated from (2). Seemingly at odds, (2) and (3) show different directions of plasticity; these can be explained via a computational model of the glomerulus taking into account the differences in stimulation protocol.

Whereas the function of GABA in olivary glomeruli has been much speculated upon since the 1970s (Llinás 1974; Sotelo et al., 1974), the excitatory terminals that are also present at those loci seem to have been largely neglected (Figure 1) (De Zeeuw et al., 1998). Therefore, it is remarkable that in this issue of *Neuron* two different labs report seminal findings regarding the effect of NMDA-dependent excitation of cells in the inferior olive (Mathy et al., 2014; Turecek et al., 2014). Stimulation of these excitatory afferents turns out to also modify the strength of olivary coupling, but the impact depends on calcium and CaMKII activation and is long-term (i.e. tens of minutes) rather than short-term (i.e. tens of milliseconds) as observed after stimulating the GABAergic afferents. The in vitro studies by Mathy et al. reveal a long-lasting decrease in the coupling-coefficient between pairs of P18-P21 olivary cells situated within 50µm from each other and subjected to 1Hz electrical stimulation of adjacent white matter. In addition, the authors show that downregulating coupling between olivary cell pairs through NMDAR activation does not depend on generation of somatic action potentials and that this activation only affects the coupling coefficient, leaving the strength of the chemical synapses unaffected. Thus, the results of Mathy et al. suggest a specific mechanism with long-lasting effects by which the coupling-coefficient between cells in the olive can be downregulated, in addition to the transient, short-term effect due to GABAR activation reported by Lefler et al. This would supply the olive with two projections using different synaptic receptor types that decrease the coupling between cells at different time scales. These findings not only underline the physiological peculiarities of the olive, but also highlight rather unique and remarkably selective mechanisms for adjusting neural connectivity.

Turecek et al. (2014) also revealed a long-lasting, NMDA and CaMKII dependent impact in vitro on the coupling-coefficient between adjacent pairs of olivary cells following stimulation of their excitatory afferents, but they employed NMDA bath solutions and electrical stimulation at 9-50Hz on tissue obtained from P24 to P50 old rats and observed an increase rather than a decrease in the strength of the coupling, while the amplitude of the subthreshold oscillations was increased. Similar enhancing effects on gap junction coupling have been reported for fish (Yang et al., 1990) and the results on oscillations are consistent with model predictions (De Gruijl et al., 2012; Torben-Nielsen et al., 2012). Upregulation of the coupling coefficient was found to be strongest in cell pairs that were only weakly coupled prior to NMDAR activation and that were spaced between 20-100µm. Using imaging techniques Turecek et al. demonstrate that the calcium influx remains highly local within the olivary dendrites. Combined with the unaltered amplitude of EPSPs from chemical synapses reported by Mathy et al., these findings suggest a highly targeted mechanism to alter the coupling coefficient between olivary neurons for longer periods of time. In fact, the two studies together offer the interesting possibility that the frequency of the excitatory input determines through the amount of calcium influx and CaMKII activation the strength of the coupling, providing a common mechanism that may allow for both up- and downregulation of coupling.

Before the short-term and long-term regulations described above can be considered as established homeostatic mechanisms, it remains to be further investigated to what extent the glomerular and extraglomerular, GABAergic terminals serve indeed different functions and to what extent a common calcium-dependent mechanism can indeed explain the differential effects upon stimulations of the NMDA-driven excitatory inputs at different frequencies. For both questions to be answered properly, it might be relevant to note that all three sets of experiments were obtained from animals at different ages; Lefler et al., Mathy et al., and Turecek et al. recorded from slices of rodents at the age of P90 to P180, P18 to P21 and P24 to P50, respectively. During development the first olivary gap junctions start to occur during the second and third week postnatally (Bourrat and Sotelo, 1983). Initially they are probably formed through assembly of both Cx45 and Cx36 hemichannels, but subsequently over the next few months they are formed solely by Cx36 hemichannels (Van Der Giessen et al., 2006). Since activation of receptors has been shown to mediate gap junction uncoupling during development (Arumugam et al., 2005), it is possible that the outcome of such processes depend on the type of connexins involved and on the precise postnatal period.

Taking aside the potential caveat of developmental issues, we ran simulations of olivary cell pairs using an extended version of a previously published multi-compartmental model (De Gruijl et al., 2012) to find out to what extent some of the seemingly contradictory findings from Lefler et al., Mathy et al. and Turecek et al. can be unified, addressing the two main questions raised above. The model, originally consisting of soma, axon hillock and main dendritic shaft compartments, was outfitted with an additional two compartments representing spine heads and high-resistance necks in an ensemble of glomeruli shared by the simulated cell pair (Figure 2A). The simulated glomeruli contained both GABAergic and NMDA receptors as well as gap junctions, and in addition we simulated the electro-diffusive properties of internal chloride and calcium concentrations inside these glomeruli, because of restricted intracellular volume (Sejnowski and Qian, 1992) (for details of model, see Supplementary Material). To assess whether glomerular and extraglomerular (i.e. dendritic), GABAergic terminals may serve different functions in controlling coupling and oscillations, we investigated whether local changes in reversal potential affect the GABAergic shunt as measured at the soma. Given the small volume of dendritic spines and the prolonged synaptic activations, we modelled the postsynaptic current at the spine head as a parallel flow of chloride and bicarbonate ions. Because of the presumptive constant bicarbonate concentration, the response to GABA application in glomeruli resulted locally in a very fast hyperpolarization, followed by a depolarization (Figure 2B). The hyperpolarization ended rapidly as the spine head accumulated chloride ions, and its driving force decreased. Despite this, the spine head potential did not plateau, because the bicarbonate had a constant gradient across the cell membrane, which caused the characteristic depolarizing waveform, reflecting synaptic conductance changes. These synaptic potentials predicted by electrodiffusion are at odds with the cell physiological observations made by Lefler et al. (2014), who argued that because of the preponderance of glomerular synapses, their contribution should determine the postsynaptic potential. To evaluate whether a combination of synaptic activation on the main dendrite and at the glomeruli could account for the recorded post-synaptic potential, we simulated low conductance dendritic synapses with a constant reversal potential in isolation and then in addition to the described above glomerular synapses (Figure 2B). Activation of synapses on the main dendrite resulted in a synaptic potential matching those reported as a somatic voltage response by Lefler et al. (2014). The addition of concurrent glomerular synapse stimulation resulted in a depolarizing shift in the steady-state shunt potential. Summation of the synaptic potential depended critically on the relative conductances, and potentially explains the reported abolition of oscillations without noticeable somatic hyperpolarization. Even though glomerular and dendritic synapses may not reflect functionally distinct ensembles, any dendritic shunt disrupts dendro-somatic current flow, which is implicated for generating oscillations (De Gruijl et al., 2012). Therefore, stimulating only glomerular synapses as well as stimulating both glomerular and dendritic synapses concurrently stop oscillations (Figure 2C and D), even in intrinsically oscillating cells.

**Figure 2.**
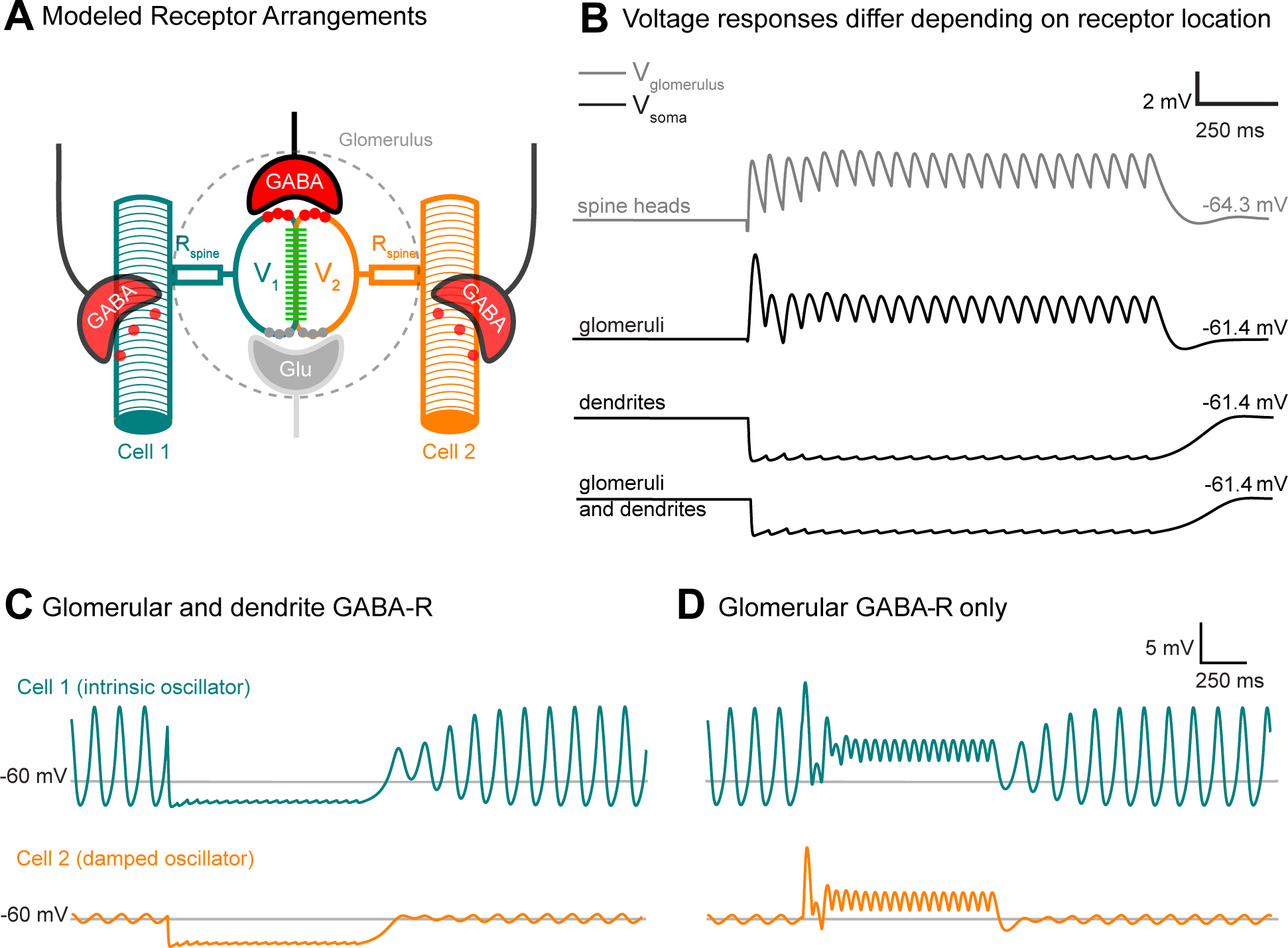
Effect of GABAergic input to glomeruli and/or dendrites of a simulated pair of coupled olivary cells. (A) Model of the glomerulus; local voltages in glomeruli of cell 1 and cell 2 (V_1_ and V_2_) are affected by GABA release in the synapse, causing Cl^-^ influx and subsequent extrusion. The local voltages V_1_ and V_2_ are further affected by the membrane potentials of their respective parent cells through highly resistive spine necks (Rspine) and gap junctional coupling (indicated in green). (B) Model predictions for stimulation patterns corresponding with 19Hz stimulation experiments conducted by Lefler et al. First trace: local response to GABA receptor activation in spine heads causes a fast hyperpolarization followed by a depolarizing response. The hyperpolarization ends due to accumulation of chloride ions in the spine head, decreasing the driving force. Due to the constant gradient of bicarbonate across the cell membrane, it is followed by a characteristic depolarizing waveform, reflecting synaptic conductance changes. Second trace: the somatic voltage during glomerular GABA activation shows a slight depolarization and reflects the response in the spine heads. Third trace: the somatic voltage during activation of GABA receptors on the main dendritic shaft causes a hyperpolarization, as the dendritic chloride concentration remains effectively unchanged due to the larger volume of the main dendritic shaft. The hyperpolarization is followed by a small calcium depolarization. Fourth trace: the somatic voltage during activation of both glomerular and dendritic GABA receptors causes the smoothest response and is depolarized with respect to the response due to exclusive stimulation of the main dendritic shaft. This response best matches the data of Lefler et al. (C) Model prediction for simultaneous dendritic and glomerular GABAergic input to a coupled cell pair. GABAergic activation temporarily abolishes oscillations without marked hyperpolarization. (D) Model prediction for exclusively glomerular GABAergic input to a coupled cell pair. GABAergic activation does not hyperpolarize the cells, but the stimulation pattern is reflected in the somatic trace.

Next, to address whether a common calcium-dependent mechanism can explain the bidirectional regulation of electrical coupling strength upon stimulations of the NMDA-driven excitatory inputs at different frequencies used in Mathy et al. and Turecek et al. we fitted the function parameters using a genetic algorithm, by optimizing gross conductance change due to their induction protocols (i.e. (I) 1Hz and (II) 25Hz synaptic stimulation paired with somatic depolarizations from Mathy et al., and (III) paired-pulse stimulation and (IV) 9Hz stimulation with no external magnesium from Turecek et al.) (Figure 3A and B). We estimated the amount of intracellular calcium in the spine head for gap junction plasticity induction protocols from both labs (without taking internal stores into account) and fitted a plasticity function (O’Donnell et al., 2011) to those results. The resulting fit was sign-consistent and could qualitatively explain both labs’ reported findings (Figure 3B). Having established that a bidirectional modulation of coupling strength between cells through NMDAR activation could be possible, we simulated a pair of olivary neurons undergoing a sudden change in coupling strength. This pair consisted of (i) an intrinsic oscillator and (ii) a damped oscillator, so as to assess the effect on both cell types. A decrease in coupling strength, as reported by Mathy et al., led to little change in the oscillation amplitude of the intrinsically oscillating cell, but caused a marked decrease in the oscillation amplitude of the damped oscillator (Figure 3C). An increase of the coupling strength, on the other hand, caused enhanced oscillations in a damped oscillator in line with the findings of Turecek et al., but again little change in the intrinsic oscillator (Figure 3D). Thus, the model’s prediction for populations of olivary cells, when also considering the observations of Turecek et al., is that the majority of cells are damped oscillators. Such cells require external input to oscillate (e.g. from coupled intrinsic oscillators), which is in line with the hypothesis that oscillations are a network effect and spread via gap junctional coupling (Torben-Nielsen et al., 2012; Lefler et al., 2014).

**Figure 3.**
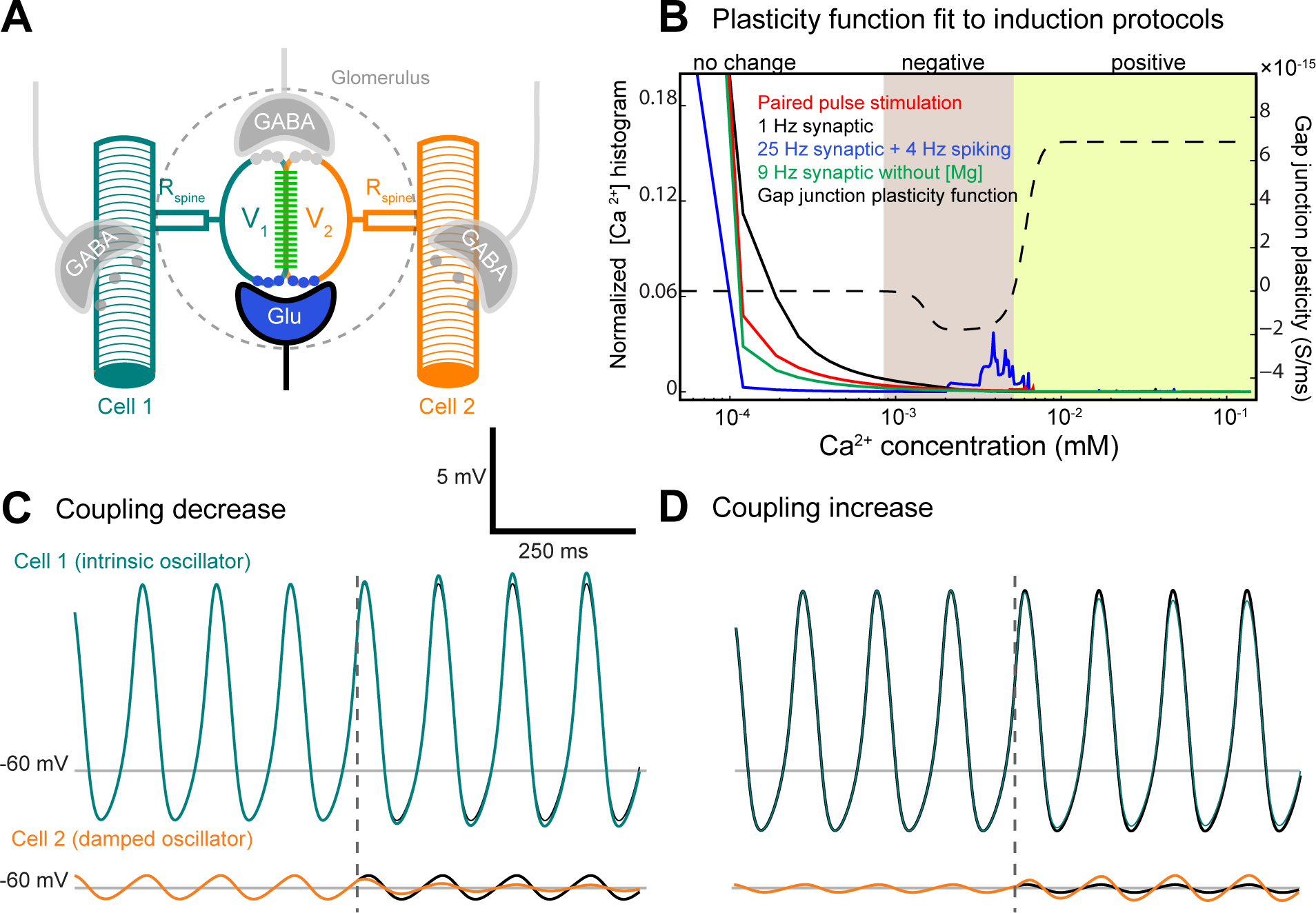
Effect of NMDAR activation on a simulated pair of coupled olivary cells. (A) Model of the glomerulus; local voltages in glomeruli of cell 1 and cell 2 (V_1_ and V_2_) are affected by NMDAR activation, causing increased intracellular Ca^2+^ concentrations. The local voltages V_1_ and V_2_ are further affected by the membrane potentials of their respective parent cells through highly resistive spine necks (Rspine) and gap junctional coupling (indicated in green). (B) Fitted curve (dashed trace) describing Ca^2+^ dependent modulation of the coupling coefficient between cell 1 and cell 2 based on results from both Mathy et al. and Turecek et al. The function shows that 1) baseline calcium concentrations cause no change (first part); 2) a small elevation of internal calcium yields a downregulation of coupling (second part, shaded in gray); and 3) a large elevation causes upregulation (third part, shaded in green). The 1 Hz synaptic activation protocol used by Mathy et al. (black trace) shows a gradual decline, with the highest probability for internal calcium concentrations in this paradigm at baseline and slightly elevated levels, causing net downregulation of coupling. The 25 Hz synaptic activation protocol used by Mathy et al. (blue trace) shows a sharp decline with some increased probabilities at higher calcium concentrations, balancing between down- and upregulation of electrical coupling. The 9 Hz stimulation paradigm as used by Turecek et al. (green trace) shows a slow decline, crosses the downregulating part of the plasticity function, ending up with a greater summed probability for upregulation of coupling. The paired pulse stimulation paradigm used by Turecek et al. (red trace) has an increased probability for higher calcium concentrations falling in the upregulating segment of the plasticity function and has a net probability for strengthening coupling. Thus, the seemingly contradicting findings of both Mathy et al. and Turecek et al. can be explained with one calcium-dependent plasticity function. (C) Model predictions for effects of coupling strength downregulation as reported by Mathy et al. (2014) on oscillations in a coupled cell pair. The effect of downregulation on an intrinsic oscillator (cell 1) is small, yielding a small increase of oscillation amplitude (reference trace in black for comparison). On the other hand, for a damped oscillator (cell 2) the reduced current input from an oscillating counterpart results in a marked decrease of oscillation amplitude (reference trace in black for comparison). (D) Effects of coupling strength upregulation as reported by Turecek et al. (2014) on oscillations in a coupled cell pair. The effect of upregulation of coupling on an intrinsic oscillator (cell 1) is small, yielding a small decrease in oscillation amplitude (reference trace in black for comparison). For a damped oscillator (cell 2), the increased current input from an oscillating counterpart serves to magnify its oscillation amplitude (reference trace in black for comparison). As the second trace fits the observations of Turecek best, it could be that most olivary neurons are damped oscillators and oscillation patterns arise in part as a network effect through Cx36 gap junctions.

The impact of different types of chemical transmission on coupling operating at different time scales open up interesting avenues for future research. It is likely that certain combinations of GABA and NMDA activity patterns have specific effects. For instance, the cerebellum may operate in an acute fashion by transiently altering inferior olive connectivity and dynamics through GABAergic feedback, overriding reflex mappings established by prior NMDAR activity. Or alternatively, coupling potentiation might be maximal through GABAR activation just prior to NMDAR activation, increasing the driving force for the calcium influx, exerting long-term effects. Follow-up experiments will surely focus on such interplay between receptor types, thereby elucidating the function and timing properties of projections to the inferior olive. In this respect it is important to note that regulation of coupling in the olivocerebellar system may play a role not only in acute motor performance, coordinating reflex-like behavior (De Zeeuw et al., 2011), but also in long-term motor learning processes (Van Der Giessen et al., 2008). The network state of the inferior olive determines the number of sodium spikes fired per event by individual cells (Maruta et al., 2007; Mathy et al., 2009) and may thereby mediate the direction and speed of learning in Purkinje cells (Mathy et al., 2009; Rasmussen et al., 2013). The phase of subthreshold oscillations in the inferior olive could be a determining factor for guiding climbing fiber induced plasticity (Mathy et al., 2009; De Gruijl et al., 2012), indicating a possible role for a GABAergic reset of olivary oscillations in both motor timing and learning. In addition, inferior olive ensemble oscillation synchrony may determine the speed and direction of cerebellar learning (Bazzigaluppi et al., 2012; De Gruijl et al., 2012), which would emphasize the importance of correct segregation of inferior olive ensembles by GABAergic input from the cerebellum. As a result, cerebellar motor execution and motor learning hypotheses are now increasingly finding common ground. Spatiotemporal firing patterns of the olivo-cerebellum affect both motor execution and plasticity, and plasticity effects take place throughout the olivocerebellar system, apparently even down to the level of electrical synapses of the inferior olive (Lefler et al., 2014; Mathy et al. 2014; Turecek et al. 2014). We may not know the exact inner workings of the olivo-cerebellar system yet, let alone that of other loci in the CNS with chemical-electrical interacting synapses, but work done in the labs of Yarom, Häusser and Welsh shows that we move towards that goal with leaps and bounds.

## References

- Arumugam, H., Liu, X., Colombo, P.J., Corriveau, R.A., Belousov, A.B., 2005. Nat. Neurosci. 8, 1720–1726.

- Bazzigaluppi, P., De Gruijl, J.R., Van Der Giessen, R.S., Khosrovani, S., De Zeeuw, C.I., De Jeu, M.T.G., 2012. Front. Neural Circuits 6.

- Best, A.R., Regehr, W.G., 2009. Neuron 62, 555–565.

- Bourrat, F., Sotelo, C., 1983. Dev. Brain Res. 8, 291–310.

- De Gruijl, J.R., Bazzigaluppi, P., de Jeu, M.T.G., De Zeeuw, C.I., 2012. PLoS Comput Biol 8, e1002814.

- De Zeeuw, C.I., Hoebeek, F.E., Bosman, L.W.J., Schonewille, M., Witter, L., Koekkoek, S.K., 2011. Nat. Rev. Neurosci. 12, 327–344.

- De Zeeuw, C.I., Holstege, J.C., Ruigrok, T.J., Voogd, J., 1990. Neuroscience 34, 645–655.

- De Zeeuw, C.I., Simpson, J.I., Hoogenraad, C.C., Galjart, N., Koekkoek, S.K.E., Ruigrok, T.J.H., 1998. Trends Neurosci 21, 391–400.

- Landisman, C.E., Connors, B.W., 2005. Science 310, 1809–1813.

- Lefler, Y., Yarom, Y., Uusisaari, M., 2014. Neuron.

- Llinas, R., 1974. Physiologist 17, 19–46.

- Llinás, R., Yarom, Y., 1981. J Physiol 315, 569–584.

- Maruta, J., Hensbroek, R.A., Simpson, J.I., 2007. J Neurosci 27, 11263–11270.

- Mathy, A., Clark, B., Häusser, M., 2014. Neuron.

- Mathy, A., Ho, S.S.N., Davie, J.T., Duguid, I.C., Clark, B.A., Häusser, M., 2009. Neuron 62, 388–399.

- O’Donnell, C., Nolan, M.F., van Rossum, M.C.W., 2011. J. Neurosci. 31, 16142–16156.

- Rasmussen, A., Jirenhed, D.-A., Zucca, R., Johansson, F., Svensson, P., Hesslow, G., 2013. J. Neurosci. 33, 13436–13440.

- Ruigrok, T.J., De Zeeuw, C.I., Voogd, J., 1990. Eur J Morphol 28, 224–239.

- Sejnowski, T., Qian, N., 1992. In: Single Neuron Computation, 117–139.

- Sotelo, C., Llinas, R., Baker, R., 1974. J. Neurophysiol. 37, 541–559.

- Spira, M.E., Yarom, Y., Parnas, I., 1976. J. Neurophysiol. 39, 882–899.

- Torben-Nielsen, B., Segev, I., Yarom, Y., 2012. PLoS Comput Biol 8, e1002580.

- Turecek, J., Yuen, G.S., Han, V.Z., Zeng, X.-H., Bayer, K.U., Welsh, J., 2014. Neuron.

- Van Der Giessen, R.S., Koekkoek, S.K., van Dorp, S., De Gruijl, J.R., Cupido, A., Khosrovani, S., Dortland, B., Wellershaus, K., Degen, J., Deuchars, J., others, 2008. Neuron 58, 599–612.

- Van Der Giessen, R.S., Maxeiner, S., French, P.J., Willecke, K., De Zeeuw, C.I., 2006. J. Comp. Neurol. 495, 173–184.

- Yang, X.-D., Korn, H., Faber, D.S., 1990. Nature 348, 542–545.

